# Spatial-temporal dynamics of contact among free-ranging domestic dogs *Canis familiaris* in rural Africa

**DOI:** 10.1101/2024.06.26.600798

**Authors:** Jared K. Wilson-Aggarwal, Cecily E.D. Goodwin, Monique Léchenne, George J.F. Swan, Metinou K. Sidouin, Matthew J. Silk, Tchonfienet Moundai, Laura Ozella, Michele Tizzoni, Ciro Cattuto, Robbie A. McDonald

## Abstract

1. Transmission of infection is affected by the spatial-temporal dynamics of host contacts. Domestic dogs *Canis familiaris* share pathogens with humans and wildlife, and managing dog-mediated diseases is a priority for public health and conservation interests.

2. We combined proximity sensors and GPS tracking to analyse spatial-temporal variation in contact among free-ranging dogs in six villages in rural Chad, during both the wet and dry seasons. We investigated dyadic interactions between dogs from different villages, the same village but different households and the same household. We assessed variation in (i) the probability of individuals having had contact, (ii) the hourly frequency of contact and (iii) contact durations.

3. Our results highlight clear seasonal and hourly patterns in contact behaviour. Contacts between dogs from different villages were rare, short in duration, primarily between male-female dyads and predominantly occurred within villages and during the dry season. Contact between dogs in the same village peaked at dawn and dusk. Sex differences were most pronounced in the wet season, where males from different households had the highest hourly contact probabilities, followed by male-female dyads. For all dogs, contact durations were longer in the dry season, but showed little hourly variation.

4. Contact patterns were not equal in space, and the probability of individuals having had contact was less than 5% when dwellings were more than 500m apart. Spatially, the probability of contact was lowest outside the village, but this increased in the dry season and peaked in the morning hours. Contact durations were notably longer outside the village, where they increased in duration for between-household dyads in the dry season.

5. At a coarse temporal scale, variation in dog contacts within and among households, and rarely between villages, may underpin seasonal variation in the incidence of dog-mediated diseases. Variation at finer temporal (hourly) and spatial scales (around households, within and outside villages) highlights the importance of routine behaviours and space use in determining patterns of contact between dogs. Practitioners should consider behavioural heterogeneities, such as those reported here, when using strategic models to support disease management decisions.

## Introduction

Interactions between individuals vary in time and space, and this can have implications for the transmission of infectious diseases (Silk et al. 2017a; Meyer & Held, 2017). Spatial-temporal patterns in contact frequency and duration between individuals are often associated with variation in the incidence of infectious diseases in both human and non-human animal hosts (Altizer et al. 2006). In humans, increased contact between children during the school term has been shown to determine broader, cyclic patterns for the incidence of measles (Bjornstad et al. 2002), chicken pox (Jackson et al. 2014) and influenza (Jackson et al. 2016). Among wildlife examples, seasonal aggregations of birds during the onset of cold weather coincide with surges in *Mycoplasma gallisepticum* infection in house finches *Carpodacus mexicanus* (Hosseini et al. 2004) and in avian influenza in water birds (Reperant et al. 2010). Associations between the dynamics of contact rates and infectious disease incidence occur because an individual’s contact behaviour can determine not only their exposure and susceptibility to infections (Drewe et al. 2011; Rimbach et al. 2015) but also their propensity to transmit diseases (Lloyd-Smith et al. 2005; Lau et al. 2017). Therefore, the capacity to forecast and control infectious diseases is dependent on our knowledge of seasonal heterogeneity in patterns of host contacts.

Free-ranging dogs *Canis familiaris* share several pathogens with humans (Otranto et al. 2017) and with wildlife (Knobel et al. 2014), and managing dog-mediated diseases is the focus of numerous public health and conservation efforts. For domestic dogs in Africa, cyclic patterns have been observed in the incidence rates of rabies (Hampson et al. 2007) and canine distemper (Viana et al. 2015), but the drivers of this remain unknown. Temporal patterns in disease incidence could be explained by multiple, interacting factors, including fluctuations in host contact rates, climatic conditions, and in the introduction of susceptible individuals (Fisman et al. 2012).

While the dynamics of dog populations are reasonably well studied (Morters et al. 2014; Conan et al. 2015), it is only in recent years that contact among free-ranging dogs has been robustly quantified, and studies have identified substantial between-individual variation in contact rates (Laager et al. 2018; Brookes et al. 2018; Wilson-Aggarwal et al. 2019; Warembourg et al. 2021). Dogs in northern Australia were shown to be highly connected with others living in their own village and the total time that each pair of dogs spent in association was typically 2-16 minutes a day (Brookes et al. 2018). Our previous, short-term analysis of free-ranging dog contact networks in rural Chad showed that individuals had heterogeneous contact rates, and that contact events were predominantly short in duration (Wilson-Aggarwal et al. 2019). Contact rates were influenced by the spatial distribution of dog-owning households and, as might be expected, dogs were more likely to interact and have longer interaction times when their households were located closer together. In one village, dogs that had larger ranges interacted with a greater number of individuals that were themselves well connected within the network (Wilson-Aggarwal et al. 2019). A study on free-ranging urban dogs in N’Djamena, Chad, found individuals were highly connected to others from nearby households, and that the spatial structure of social communities was characterised by elements of urban infrastructure (e.g. main roads) that restricted dog movements (Laager et al. 2018). Finally, a comparative study highlighted that, while the structure of dog contact networks were similar, and the spatial distribution of households was important in determining contact, other predictors (including individual and household traits) were highly variable within and between countries; Chad, Guatemala, Indonesia and Uganda (Warembourg et al. 2021).

These studies have each provided a ‘snapshot’ account for the contact behaviour of free-ranging dogs, over relatively short observation periods (3-10 days). Therefore, it is not known whether or how contact events are subject to temporal or spatial variations in occurrence or duration. Our previous work on the ranging behaviour of dogs in rural Chad revealed periodicities in activity and space use, hourly variations in the propensity for dogs to be “at home”, and showed that dogs ranged further during the dry season when there is little rain and vegetation (Wilson-Aggarwal et al. 2021). Given space use determines opportunities for contact, we might also expect contact probabilities to vary throughout the day, and for contacts between dogs in different households and villages to be higher in the dry season. Such patterns of contact can be important in determining the trajectories of disease epidemics (Volz & Meyers, 2009; Enright & Kao, 2018). For example, simulations of rabies in racoons *Procyon lotor*, showed seasonal outbreaks were best explained by shifts in contact durations, rather than observed shifts in interactions between sexes or pulses in birth rates (Hirsch et al. 2016). Evidence suggests that the temporal dynamics of contacts have larger influences on transmission dynamics when R_0_ (the mean number of secondary cases of an infected individual in a fully susceptible population) is <2 (Chen et al. 2014), or when the disease shows low transmissibility and a long infectious period (Springer et al. 2017). Knowing how the contact rates of dogs vary in time and space might therefore be especially important for understanding the dynamics of such diseases in dogs, including rabies (Hampson et al. 2009; Kurosawa et al. 2017) and canine distemper (Viana et al. 2015).

In this study, we investigated spatial-temporal variations in contacts among free-ranging domestic dogs in six rural villages in Chad, where rabies and other dog-mediated zoonoses remain endemic. Specifically, we set out to test the hypotheses that the contact behaviour between free-ranging dogs and the duration of their contacts (1) are seasonally variable and greater in the dry season when dogs range further, (2) are spatially variable and greater within the village, where dogs are more likely to encounter one another and, (3) show fine-scale temporal (hourly) patterns reflective of periodicity in their ranging behaviour.

## Methods

### Field study timings and locations

Fieldwork was conducted in six rural villages in Chad (Figure 1) during the dry season (between 5^th^ March and 17^th^ May 2018) and again during the wet season (between 3^rd^ August and 17^th^ October 2018). The most northerly village, Medegue (11°01’48.8“N 15°26’37.7”E), is located on a main road 15 km from the district town of Guelendeng in the Mayo-Kebbi East region of Chad. The remaining villages are in the Moyen-Chari region, and can be split into two geographical sites, ‘Sarh East’ and ‘Sarh West’. Sarh West is 10 km to the west of Sarh, the provincial capital, and includes: Ngakedji (9°11’16.5“N 18°18’10.7”E); Kira (9°10’50.8“N 18°17’00.3”E) which is ∼2km from Ngakedji; and Bembaya (9°11’33.6“N 18°17’42.3”E) which is between the two villages. Sarh East is 40km to the east of Sarh town and includes: Marabodokouya (9°19’42.3“N 18°43’20.0”E), a large village that is split into hamlets up to 10km apart; and Tarako, a small, nucleated hamlet of the village Tarangara (9°08’19.8“N 18°42’00.9”E), that is ∼20km south of Marabodokouya’s village centre.

**Figure 1.**
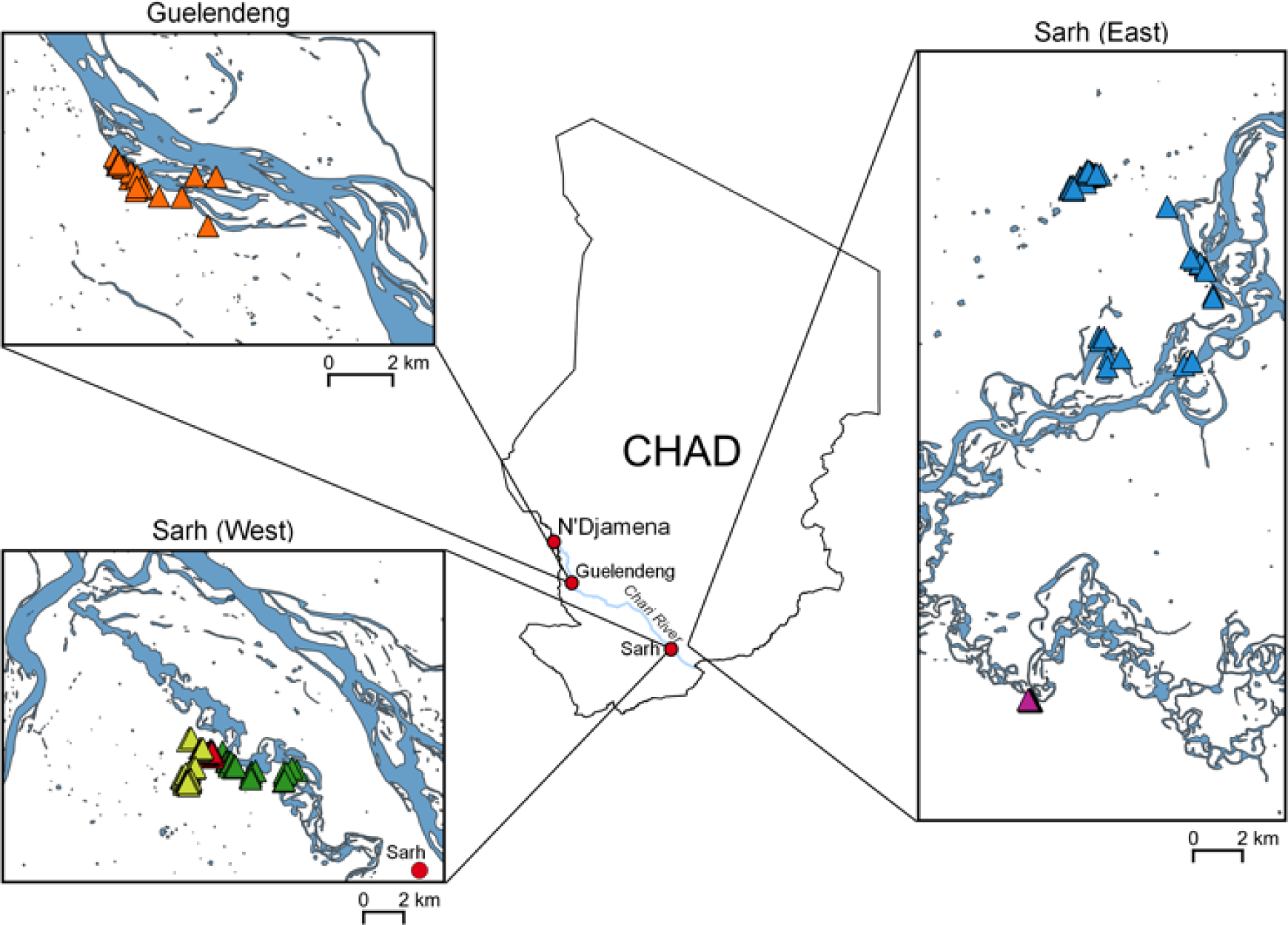
Locations in Chad of six study villages and the households where dogs were collared. Triangles represent dog-owning households. In Guelendeng, orange triangles represent households from the village Medegue. In Sarh (East), blue triangles represent households in the village of Marabodokouya and pink triangles are households in Tarangara. In Sarh (West), yellow triangles represent the village Kira, green triangles the village Ngakedji and red triangles represent Bembaya. The locations of the capital city N’Djamena, Guelendeng and Sarh are shown for reference.

In all settlements, the households were not fenced and the dogs were kept outside, where they were free to roam. With the verbal consent of dog owners and the village chief, dogs were collared with standard nylon dog collars (Ancol Heritage). We recorded the dogs’ sex and age in months (as recalled by the owner). All dogs were sexually intact, had clear ownership and were closely affiliated with a household. Puppies (< 6 months of age) were not collared. We recorded the location of the collared dogs’ households using a handheld GPS (Garmin GPSMap 64S). Since we visited all households known to have dogs, the number of dogs owned by each household was recorded and summed to estimate the adult dog population. Collars were fitted with (1) an i-GotU GT-600 GPS unit (Mobile Action Technology Inc., Taiwan) configured with a fix interval of 10 minutes and, (2) a proximity sensor developed by the OpenBeacon project (http://www.openbeacon.org/) and the SocioPatterns collaboration consortium (http://www.sociopatterns.org/). The proximity sensors exchange several radio packets per second and proximity is measured by the attenuation, defined as the difference between the received and transmitted power (Cattuto et al. 2010; Isella et al. 2011; Wilson-Aggarwal et al. 2019). An attenuation threshold of −70dbm was used, as it has been shown to identify close-contact events (within 1–1.5m) during which a communicable disease might be transmitted (Stehle et al. 2013; Voirin et al. 2015; Wilson-Aggarwal et al. 2019).

### Data processing

Processing of the proximity data was conducted in Python v2.7. Data were cleaned by assessing inter-logger variability and comparing the number of packets emitted and received for all pairs of sensors. The data were discarded if they showed deviations from the expected linear relationship between radio packets emitted and received. A contact event was defined by sensors exchanging radio packets for a minimum of 20 consecutive seconds. The contact event ended when the sensors stopped exchanging radio packets in any of the subsequent 20s periods.

Processing of GPS data was conducted in R v3.3.3 (R Core Team, 2017) using the most appropriate coordinate reference system for Chad (EPSG:32634). The distance between households for each pair of individuals (dyad) was calculated. GPS data were cleaned by removing erroneous fixes implying speeds greater than 20km/hr between consecutive fixes. To give time for dogs to adjust to the collars, we discarded GPS and proximity records 12 hours after collar deployment and 12 hours before collar collection.

### Locating contacts

To identify the locations where contact events were first initiated, continuous time movement models were fitted to the spatial data using the ‘ctmm’ R package (v0.5.5). The models were used to simulate the possible location of individuals at times between fixes; For each individual, 30 realisations of their possible paths were simulated at an interval of one minute. For the start time of each contact event, the mean latitude and longitude of simulated paths were calculated and taken as the location where contact was most likely to have been initiated. Contact locations were categorised as either: around the household (within 50m of either dogs’ household), in the village (>50m of either dogs’ household but within 100m of any household in the village that owned a tracked dog), or outside the village (>100m from any household in the village that owned a tracked dog).

For each dyad, we recorded whether or not the individuals had contact (0/1), how many contacts they had and the duration of each contact event. We then calculated the daily probability of contact for each dyad; number of days a contact was recorded divided by the number of days the individuals were contemporaneously observed. In addition, the daily frequency of contact for dyads at each hour of the day and location category was calculated; the number of days at least one contact was recorded at hour *i* and at location category *j*, divided by the number of days a contact could have been recorded at hour *i*. Finally, for each dyad and day observed, the total duration of contact events initiated in hour *i* was calculated.

### Statistical analysis

Generalised linear mixed-effect models (GLMMs) were used to investigate variation in contact at the dyad level (Cross et al. 2012; Cross et al. 2013; Silk et al. 2017b). Preliminary analysis suggested that contacts between dogs from different villages were relatively rare (see results), and so models were only run for within-village dyads, with separate models for within- and between-household dyads.

To investigate predictors of whether contact between dogs in a dyad had been recorded (0/1), a GLMM was fitted with a binomial error structure using the R package ‘lme4’ (v1.1-18.1; Bates et al. 2015). Explanatory variables included the dyads sex composition (categorised as male-male, male-female or female-female), age difference in months, the log (base 2) distance between the individuals’ households for between-household dyads, season, and the two-way interactions between these variables and season. The log (base 10) number of minutes that dyads were contemporaneously observed was included as a fixed effect to control for variation in observation times. The identity of each individual in a dyad and the identity of each household were included as random effects, to account for repeated observations.

To investigate variations in the daily frequency of contact at each hour of the day and location for dyads where contact had been recorded, a GLMM was fitted with a binomial error structure. Because the response variable was a proportion, hours of the day for dyads that had a low sample size (<10 days contemporaneously observed) were removed from the analysis. Fixed effects included those described above, except for minutes observed, and with the addition of the hour of day (as a categorical variable), location, and the two-way interactions of these two variables with season. Random effects were the same as in the previous model, with the addition of dyad ID, to account for repeated observations.

To investigate spatial-temporal variation in the duration of contact events, a GLMM was fitted with a negative binomial error structure using the R package ‘glmmTMB’ (v0.2.3). To ensure models converged, data on dyads from Tarangara village were excluded as data from the wet season were absent due to flooding preventing our access to the village. Fixed and random effects were the same as the previously described model.

The R package ‘emmeans’ (v1.3.3) was used to estimate test statistics, *p*-values, odds ratios and confidence intervals for contrasts from the full model. The R package ‘DHARMa’ (v0.2.6) was used to assess model residuals and goodness of fit. The marginal R^2^ (proportion of total variance explained by fixed effects) and conditional R^2^ (proportion of total variance explained by fixed and random effects) were extracted from the models. Full model results are reported in the supplementary material, and only statistically significant findings are reported in the results section.

## Results

In total we visited 161 households that on average had 1.6 dogs (range = 1-7 dogs). Dogs were collared for a mean of 37 days (1–70 days) in the dry season and 34 days (2–65 days) in the wet season. Proximity data were available for 199 and 166 individuals in the dry and wet seasons respectively and, of these, we collected spatial data for 179 and 149 individuals.

### Between-village contacts

During the dry season in Sarh East, none of the 1400 between-village dyads were recorded to have been in contact. In Sarh West, 25 (1%) of the 2013 between-village dyads came into contact during the dry season, with 166 contact events recorded and a median contact duration of 20s (20–40s; Table S1). Between-village contact events were most frequently between male-female dyads (n = 139, 82%). Contacts tended to occur either at 6am or between 7-8pm (Figure S3). Of the locatable contact events, most (n = 101, 63%) occurred in the village, 43 (27%) occurred around the household and 16 (10%) occurred outside the village. One male dog accounted for 105 (63%) contact events, which occurred over 1 week and involved 6 dogs living in 5 households located 500-900m from its own household. Similarly, one female dog accounted for a further 44 (27%) contact events, which occurred over 2 weeks and involved 5 dogs from 4 households. Very occasional (n = 4) longer distance contact events were recorded between dogs living in households 1600-2400m apart.

In the wet season, only 3 of 1556 (<1%) between-village dyads in Sarh West came into contact. A total of 5 contact events were recorded, all of which were 20s long and located in the village. These events involved 2 dogs from the same household that, over the course of 2 days, encountered 3 dogs from different households located 300-500m from their household.

### Within-village contacts

In the dry season, 713 of 4254 dyads (17%) were recorded to have had contact (Table 1). 47,373 contact events were recorded, of which 34,890 (74%) were for within-household dyads and 12,483 (26%) for between-household dyads. For within-household dyads, median daily probability of contact was 27% (inter-quartile range = 18–32%) and median total duration of contact was 100s (40– 260s; Figure S2). For between-household dyads, the median daily probability of contact was 5% (2– 10%) and the median total duration of contact was 60s (20–120s).

**Table 1.** Summary of recorded contacts among free-ranging domestic dogs from six villages in rural Chad. For each season and village, the adult dog population size is reported along with the number and proportion of individuals sampled, the number of observable within-village dyads, the number of recorded within-village dyads recorded to have had contact, and the number of contact events. The percentage for the number of dyads observed in contact is calculated using the population count for villages. For within- and between-household dyads that had recorded contact, the median and inter-quartile range are reported for the daily probability of contact (number of contacts divided by the number of days monitored). For all dyads in contact, the median and inter-quartile range are provided for the proportions of contact events that were around the household, in the village and outside of the village. In Tarangara village, no dogs were collared in the wet season.

| Village | Season | Pop. | Individuals sampled | No. of observable dyads | No. of dyads in contact | No. of contacts | Within household daily probability (%) | Between household daily probability (%) | Proportion (%) around the household | Proportion (%) in the village | Prop. outside the village (%) |
| --- | --- | --- | --- | --- | --- | --- | --- | --- | --- | --- | --- |
| Medegue | Dry | 30 | 26 (87%) | 325 (75%) | 62 (19%) | 3814 | 20 (18-23) | 6 (3-10) | 0 (0-36) | 51 (28-76) | 11 (0-38) |
|  | Wet | 38 | 29 (76%) | 355 (50%) | 61 (17%) | 4401 | 51 (42-62) | 5 (3-11) | 0 (0-12) | 57 (48-100) | 1 (0-47) |
| Kira | Dry | 45 | 44 (98%) | 942 (95%) | 168 (18%) | 15015 | 32 (20-58) | 4 (2-10) | 0 (0-32) | 86 (41-100) | 0 (0-6) |
|  | Wet | 49 | 36 (73%) | 618 (53%) | 210 (34%) | 18529 | 69 (42-84) | 7 (3-17) | 0 (0-18) | 93 (58-100) | 0 (0-7) |
| Bembaya | Dry | 12 | 12 (100%) | 65 (98%) | 46 (71%) | 5586 | 24 (19-73) | 8 (4-21) | 20 (1-48) | 55 (24-78) | 0 (0-30) |
|  | Wet | 21 | 18 (86%) | 151 (72%) | 85 (56%) | 4535 | 45 (21-63) | 14 (5-26) | 0 (0-25) | 86 (49-100) | 0 (0-4) |
| Ngakedji | Dry | 27 | 27 (100%) | 341 (97%) | 47 (14%) | 4499 | 16 (11-17) | 9 (4-17) | 6 (0-35) | 65 (34-100) | 0 (0-21) |
|  | Wet | 25 | 17 (68%) | 136 (45%) | 9 (7%) | 649 | 14 (9-18) | 7 (4-17) | 50 (48-72) | 29 (24-42) | 0 (0-10) |
| Marabodokouya | Dry | 85 | 70 (82%) | 2391 (67%) | 347 (15%) | 15693 | 29 (24-34) | 5 (2-9) | 0 (0-22) | 14 (0-60) | 60 (0-100) |
|  | Wet | 111 | 66 (59%) | 1961 (32%) | 167 (9%) | 5145 | 32 (12-68) | 10 (3-18) | 0 (0-46) | 50 (0-100) | 0 (0-47) |
| Tarangara | Dry | 21 | 20 (95%) | 190 (90%) | 43 (23%) | 2766 | 8 (5-19) | 5 (3-11) | 48 (20-91) | 48 (0-75) | 0 (0-0) |
|  | Wet | - | - | - | - | - | - | - | - | - | - |
| Overall | Dry | 220 | 199 (91%) | 4254 (76%) | 713 (17%) | 47373 | 27 (18-32) | 5 (2-10) | 0 (0-33) | 44 (0-100) | 4 (0-100) |
|  | Wet | 244 | 166 (68%) | 3221 (38%) | 533 (34%) | 33259 | 46 (21-71) | 8 (3-18) | 0 (0-25) | 75 (36-100) | 0 (0-18) |

In the wet season, 533 of 3221 dyads (17%) were observed to have had contact. 33,259 contact events were recorded, of which 24,793 (75%) were for within-household dyads and 8,466 (25%) were for between-households dyads. For within-household dyads, median daily probability of contact was 46% (21–71%) and median total duration of contact was 100s (40–280s). For between-household dyads, median daily probability of contact was 8% (3–18%) and median total duration of contact was 20s (20–100s).

### Likelihood of having had contact

The odds of contact having been observed for between-household dyads reduced by a factor of 0.36 for every doubling in the distance between the dogs’ households (95% confidence interval (CI) = 0.33–0.40, z = −19.06; p < 0.001; Figure 2). Compared to in the dry season, the odds of contact for male-male dyads was 2.78 (CI = 1.52–5.10; z = 3.31; p = 0.001) times higher in the wet season, where the odds of contact for male-male dyads was also 1.74 (CI = 1.07–2.82; z = 2.68; p = 0.020) times higher than for male-female dyads and 2.52 (CI = 1.18–5.37; z = 2.90; p = 0.012) times higher than for female-female dyads. In the dry season, contact between male-female dyads was 1.60 (CI = 1.02– 2.52; z = 2.44; p = 0.039) times higher than female-female dyads. The odds of a dyad having had contact increased by a factor of 2.57 (CI = 1.55–4.26; z = 3.64; p < 0.001; Figure S2) for every 10-fold increase in the number of minutes observed.

**Figure 2.**
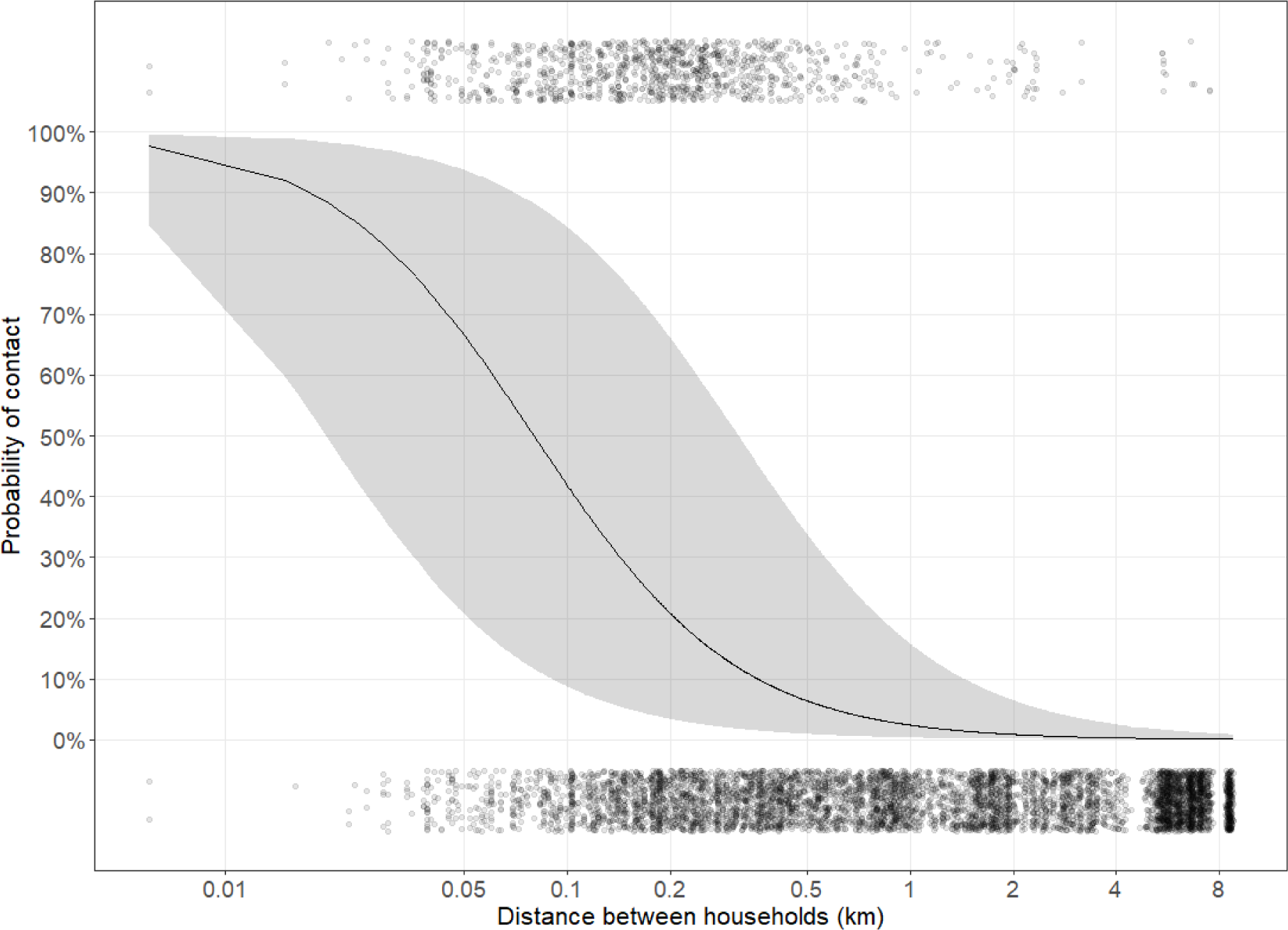
The probability of contact between dyads of free-ranging domestic dogs from different households in rural Chad, with increasing distance between the dogs’ households. The predictions and confidence intervals are plotted from a general linear mixed model and conditioned on the random terms. The points are the raw data and have a vertical jitter applied. The x axis is on a log_2_ scale.

No significant fixed effects were identified in the model for the odds of contact between dogs living within the same household (Table S2).

### Contact frequency; between-household dyads

For dogs living in different households and that were observed to have had contact, the odds of contact were 0.79 (CI = 0.70-0.89; z = −3.88; p < 0.001) times lower in the dry season than in the wet season. An interaction between season and the distance between households was identified (z = - 3.18; p = 0.002), whereby the odds of contact reduced with every doubling in the distance between the dogs’ households by a factor of 0.61 (CI = 0.57–0.66) in the dry season and 0.68 (CI = 0.63–0.74) in the wet season. During the dry season, contact events peaked between 5-7am and again between 6-8pm, while a peak was only observed between 6-8pm in the wet season (Figure 3A; Table S3).

**Figure 3.**
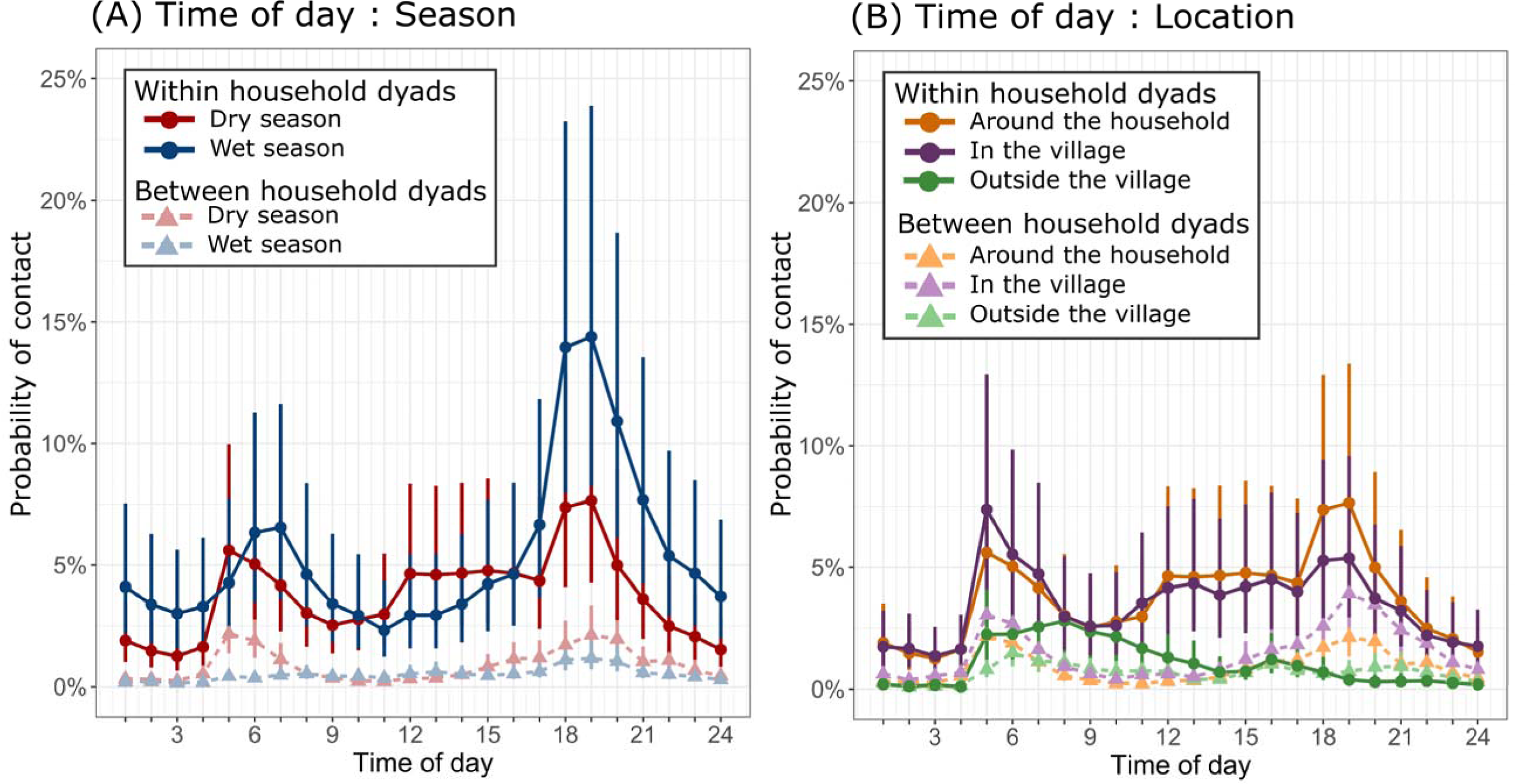
Spatial-temporal variation in the hourly probability of contact events among free-ranging domestic dogs in rural Chad. Predictions and confidence intervals are derived from general linear mixed models. A) Predicted hourly probability of contact between dogs in the dry (red) and wet (blue) seasons. B) Predicted hourly probability of contact between dogs in different locations; around the household (orange), in the village (purple) and outside of the village (green). Results for within-household dyads are illustrated with bold circles, and between-household dyads as light triangles. In both plots, the predictions for the between-household dyads were calculated with the distance between households set to the lowest observed value.

Compared to the dry season, the odds of contact in the wet season was 1.81 (CI = 1.43–2.30; z = 4.88, p < 0.001) times higher for male-male dyads and 1.20 (CI = 1.06–1.35; z = 2.92, p = 0.004) times higher for male-female dyads. Within season differences in the mixing among/between sexes were only evident in the wet season such that, the odds of contact for male-male dyads was 1.58 (CI = 1.16-2.16; z = 3.44; p = 0.002) times higher than for male-female dyads and 2.38 (CI = 1.52-3.72; z = 4.51; p <0.001) times higher than for female-female dyads, while the odds of contact was 1.50 (CI = 1.09 – 2.07; z = 3.01; p = 0.008) times higher for male-female dyads than female-female dyads.

Contact behaviour at different locations varied by season whereby, compared to in the wet season, the odds of contact in the dry season was 1.67 (CI = 1.42–1.97; z = 6.18; p < 0.001) times higher outside the village, 0.63 (CI = 0.55–0.73; z = −6.53; p < 0.001) times lower around the household and 0.47 (CI = 0.41–0.53; z = −12.17; p < 0.001) times lower in the village. Within season differences in the odds of contact between locations was consistent but most exaggerated in the wet season, whereby the odds of contact in the village was 2.30 (CI = 2.09-2.52; z = 20.64; p < 0.001) times higher than around the dogs’ households, 7.35 (CI = 6.36-8.50; z = 32.20; p < 0.001) times higher than outside the village, and 3.20 (CI = 2.74-3.74; z = 17.50; p < 0.001) times higher around the dogs’ households than outside the village. The location of contacts also varied with time of day (Figure 2B; Table S3), and contact events outside the village had a single, small peak at 6am, followed by a gradual decline over the rest of the day. Contact events in the village and around the households peaked at 6am and again at 7pm.

### Contact frequency; within-household dyads

For dogs living in the same household and that were observed to have had contact, the odds of contact was 1.14 (CI = 1.03-1.26; z = 2.46; p = 0.014) times higher in the wet season than in the dry season. Contact events peaked between 5-7am and 6-7pm (Figure 3A), with a more pronounced evening peak in the wet season. Differences in the sex composition of dyads varied between but not within seasons, whereby the odds of contact in the wet season was 1.93 (CI = 1.72–2.16; z = 11.45; p < 0.001) times higher for female-female dyads than in the dry season, and 0.75 (CI = 0.58–0.98; z = - 2.08; p = 0.038) times lower for male-male dyads.

Contact frequency at each location category varied between seasons such that, compared to in the dry season, the odds of contact in the wet season was 0.82 times lower around households (CI = 0.74-0.91; z = −3.59; p < 0.001) and 1.82 times higher in the village (CI = 1.64-2.02; z = 11.22; p < 0.001). Within season differences in the odds of contact between location categories were also evident and, in the dry season, the odds of contact outside the village was 0.20 (CI = 0.18-0.22; z = 38.78; p < 0.001) times lower than in the village and 0.19 (CI = 0.17-0.21; z = 39.78; p < 0.001) times lower than around the household. During the wet season, the odds of contact in the village was 9.34 (CI = 8.32-10.48; z = 45.28; p < 0.001) times higher than outside the village and 2.12 (CI = 1.99–2.28; z = 25.83; p < 0.001) times higher than around the household, while the odds of contact around the household was 4.39 (CI = 3.89-4.96; z = 28.64; p < 0.001) times higher than outside the village. The temporal pattern of contact probabilities throughout the day and for different locations was similar to that of the between-household dyads (Figure 3B).

### Contact durations; between-household dyads

For dogs living in different households, the duration of contact events decreased by a factor of 0.97 (CI = 0.93–1.00; z = −2.15; p = 0.032) for every doubling in the distance between households. The duration of contact events recorded outside the village was 1.45 (CI = 1.26–1.68; t = 5.12; p < 0.001; Table S3; Figure 4A) times longer in the dry season than in the wet season. During the dry season, contact durations were 1.35 (CI = 1.21–1.52; t = 6.28; p < 0.001) times longer outside the village than in the village, and 1.38 (CI = 1.22–1.57; t = 6.01; p < 0.001) times longer than around the household. The duration of contacts varied with time of day and location, with shorter contact durations outside the village after 5pm, but longer durations around the household and in the village at 5am (Figure 4B). The duration of contact events increased by a factor of 1.01 (CI = 1.00–1.01; z = −2.43; p = 0.015) for every 1 month increase in age differential between dogs.

**Figure 4.**
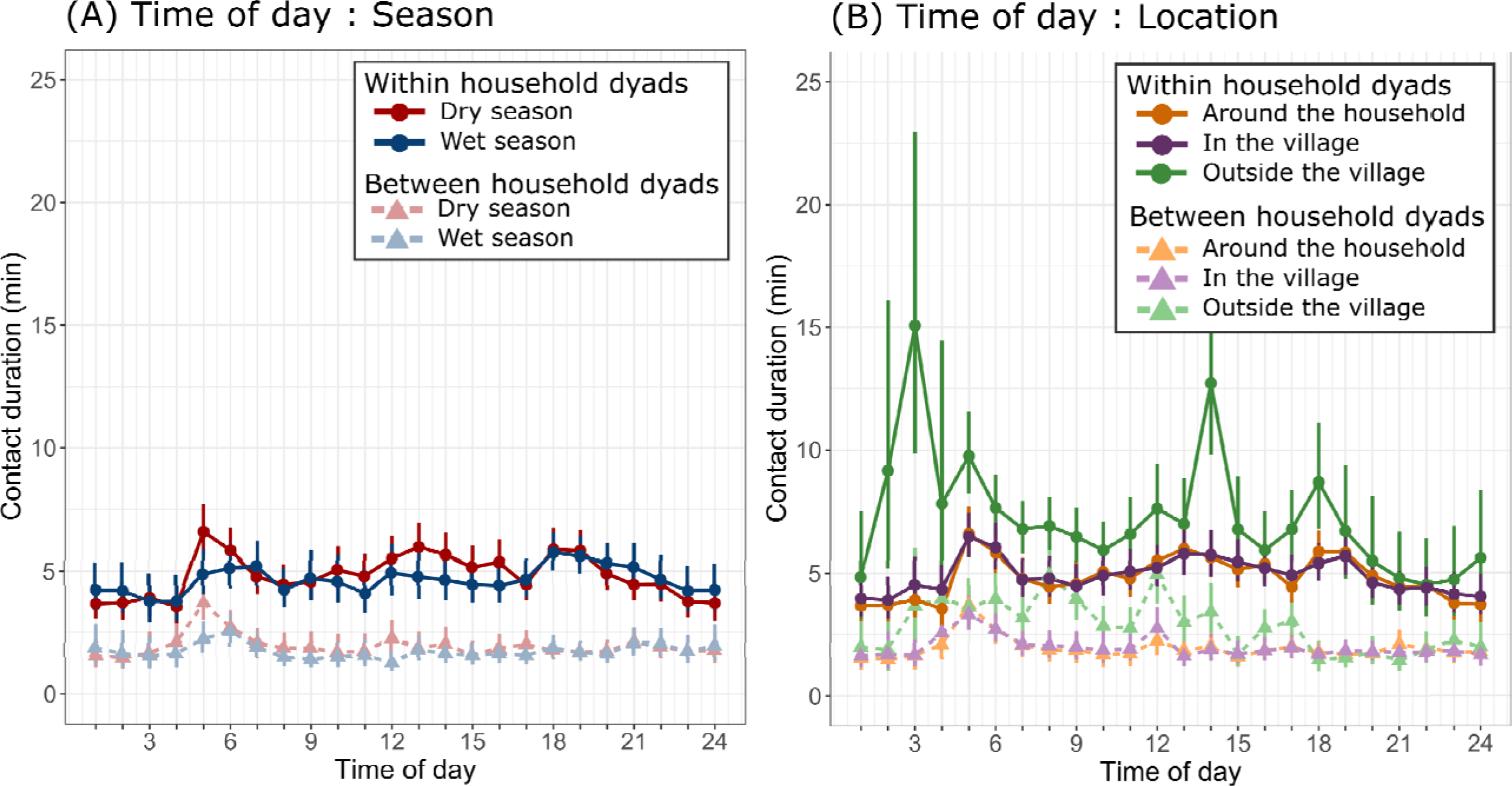
Spatial-temporal variation in the hourly duration of recorded contact among free-ranging domestic dogs in rural Chad. Predictions and confidence intervals are derived from general linear mixed models. A) Predicted durations of contact between dogs in the dry (red) and wet (blue) seasons, B) Predicted durations of contact in different locations: around the household (orange), in the village (purple) and outside of the village (green). Results for within-household dyads are illustrated with dark circles and between-household dyads as light triangles. In both plots, the predictions for the between-household dyads were calculated with the distance between households set to the lowest observed value.

### Contact durations; within-household dyads

For dogs living in the same household that were observed to have had contact, the duration of contact events outside the village was 1.46 (CI = 1.31–1.64; t = 6.58; p < 0.001) times longer in the dry season than in the wet season. During the dry season, the duration of contacts outside of the village was 1.42 (CI = 1.29–1.57; t = 8.42; p < 0.001) times longer than those in the village and 1.46 (CI = 1.32–1.61; t = 8.83; p < 0.001) times longer than those around the household. While there were significant differences in the duration of contacts at different times of the day, there was no obvious pattern or notable differences between the seasons (Figure 4A; Table S3). The duration of contacts varied most notably for contacts outside the village, where peaks occurred at 3am and 2pm (Figure 4B). The marginal R^2^ (0.02) and conditional R^2^ (0.03) imply the fixed and random effects provide poor predictive power.

## Discussion

We have provided an in-depth account of the spatial-temporal contact patterns for free-ranging domestic dogs in multiple villages in rural Chad. We found that while variation in contact behaviour is somewhat determined by the distribution of households and villages, there was marked seasonal and hourly variation in the likelihood, frequency and duration of contact events between dogs. Moreover, contact behaviour varied spatially and we found evidence for seasonal differences in the preferential mixing between sexes.

The observed temporal patterns of contact between dogs complement the seasonal and hourly variations in their ranging behaviour (Wilson-Aggarwal et al. 2021). Dogs living in the same village had a greater likelihood of contact in the wet season, when individuals had smaller home ranges that were more concentrated around the household. By contrast, occurrence of the rarer contacts for between-village dyads increased in the dry season, when dogs’ home ranges greatly increased.

Hourly contact frequencies varied by location, with a peak in the morning hours for contacts outside the village, and peaks around the household and within the village at dawn and dusk, the timings of which vary little in Chad. The daily periodicity of the dogs’ space use and activity levels (Wilson-Aggarwal et al. 2021) can explain variation in hourly contact rates, whereby contact occurs when individuals pass other dog-owning households as they range around and beyond the village in the morning, and when they return home in the evening.

The spatial-temporal variation in contacts among dogs will influence the likelihood of introduction and onward transmission of infections and, thereby, variation in seasonal risks and trajectories of disease outbreaks. In terms of rabies, dog infections in Africa persist all year, with monthly fluctuations in incidence rates (Koeppel et al. 2021; Hikufe et al. 2019; Kidane et al. 2016; Munang’andu et al. 2011; Ali et al. 2010). Incidence rates for rabies in rural Chad are currently not available, however, as seen in Namibia (Hikufe et al. 2019) and Ethiopia (Ali et al. 2010), we might expect cases to increase locally during the wet season, when dogs’ home ranges are smaller and between-household contacts are more frequent. By contrast, cases of rabies in dogs might become more spatially widespread in the dry season, with more opportunities for transmission to other villages, due to dogs ranging further and having higher probabilities of contact with individuals from different villages. On their own, the contact patterns observed here cannot explain the 3-6 year cycles of rabies described for free-ranging dog populations in Africa, which are likely due to a combination of factors such as reactive vaccination campaigns and the high turnover of dog populations (Hampson et al. 2007). However, this structured variation in contact rates should be incorporated into models of disease transmission to assess the role of such variation in disease dynamics, and to support improved forecasting of outbreaks and the identification of novel management strategies.

The impact of the dyads sex composition on contact rates and duration also differed with respect to season. Whilst we found no seasonal differences in the likelihood of recording contact between males and females, if contact had been recorded, the hourly frequency of contact for between-household dyads was greater in the wet season than in the dry. Similarly, contact in male-male between-household dyads were more likely to be recorded and had higher hourly contact frequencies in the wet season. Although we could not determine the nature of contact events, aggressive interactions have been shown to be more common between male dogs (Pal, 2015), and this may increase the risk of disease transmission through behaviours such as biting. Free-ranging domestic dogs are generally thought of as non-seasonal breeders (Lord et al. 2013), and evidence for seasonal breeding has only been found for dogs in India (Pal, 2011; Fielding et al. 2021). The populations in this study were unmanaged and reproductively intact, and seasonal mixing between the sexes could be indicative of seasonality in some aspects of breeding behaviour, in this case likely stemming from variation in climate and mobility.

Contact events between dogs are predominantly brief interactions and, as in our earlier study in Chad (Wilson-Aggarwal et al. 2019), generally lasted 60s or less. However, we found that contact duration depended on where and when events took place; in the dry season contact events initiated outside of the village were longer than those around the household or in the village. The spatial and temporal context for contact in this population is therefore material to the nature of interactions.

Contact frequency and duration can both determine whether infection is transmitted (Smieszek, 2009), and the importance of each depends on the infectiousness of the disease and its mode of transmission (Cao et al. 2014; Toth et al. 2015). Therefore, the transmission of diseases requiring prolonged direct contact may be increased when dogs encounter one and other outside of the village bounds.

A network analytical approach could provide a means of identifying traits associated with the position of individuals/households within the contact network (as determined by measures of centrality). This would, however, require more synchronous observation of individuals, which we were unable to achieve in the wet season due to logistical constraints. Another further opportunity would be to describe in greater detail the proximate drivers of variation in dog contacts and their contact behaviour, and the degree to which these are dog-or owner-mediated. This distinction could be important for the management of diseases, since owner-mediated behaviours imply a greater opportunity for owners to manage contact between dogs, and household level traits related to husbandry could provide simple guidance for targeting management strategies, which aim to prioritise individuals that present greater epidemiological risks.

Our results can inform the use of frequency-dependent functions such as spatial and/or social scaling parameters used in disease models to generate variation in dog contact rates (Beyer et al. 2011; Johnstone-Robertson et al. 2017). For example, our results can help specify such scaling functions to better simulate within-patch dynamics of metapopulation models, which have been used to highlight how long-range human-mediated movements of dogs facilitate the persistence of rabies in Central African Republic (Colombi et al. 2020). Scaling the probability of contact between dogs using the distance between their households could adequately capture the spatial variation in contact, including rare between-village contacts. For free-ranging dogs in rural Chad this would involve a shape function that reduces the probability of contact between dogs from different households to below 5% when their households are greater than 500m apart. Seasonal mixing patterns between the sexes and marked variation in the likelihood and frequency of contacts among individuals and households (as expressed through the models’ random terms), suggest a social scaling parameter is also required to accurately simulate contact patterns.

Our study provides a thorough account of the spatial-temporal variation in contact among free-ranging domestic dogs, living in close affiliation with people in rural Chad, where this species shares several pathogens with humans and wildlife (McDonald et al. 2020; Goodwin et al. 2022; Naïssengar et al. 2021). The clear seasonal variation we identify in the likelihood, duration and location of contact events have implications for the potential management of disease risk, controlling outbreaks and for epidemiological models of dog-mediated diseases. By quantifying the interactions of these domestic animals in the contexts of space, as structured by human households, and time, primarily structured by seasonal variation, these insights highlight the merits of a One Health approach to improving human and non-human animal health.

## Acknowledgments

JWA was funded by a Natural Environment Research Council studentship from the GW4 + Doctoral Training Partnership and by the University of Exeter. The Carter Center, World Health Organization (WHO), and Chad Ministry of Public Health funded fieldwork and provided in-country support for work on dogs, as part of a separate project on Guinea worm epidemiology. We thank Phillip Tchindebet Ouakou, Mario Romero, Ernesto Ruiz-Tiben, Adam Weiss, Hubert Zirimwabagabo, and all the field and operations staff of The Carter Center, WHO, and Chad Ministry of Public Health, and local government officials for their advice and support during this project. We also thank Richard Ngandolo and Fayiz Abakar from the Institut de Recherche en Elevage pour le Développement for facilitating importation of project materials. This work was approved by the ethics committee of the College of Life and Environmental Sciences (Penryn Campus) of the University of Exeter (ref 2018/2318) and was facilitated by the Chad Ministry of Public Health as part of a related program of work on the epidemiology and ecology of Guinea worm in dogs. CC, LO, and MT acknowledge partial support from the Lagrange Project of the ISI Foundation funded by CRT Foundation.

## Author contributions

RM and JWA conceived the study. Data collection was conducted by JWA, CEDG, GJFS, ML, MKS and TM, with technical support from LO, MT and CC. Data analysis was done by JWA with advice from MJS, LO and MT. JWA and RM wrote the original draft and all authors reviewed and contributed critically to the final manuscript.

## Data Availability Statement

Data will be deposited on dryad once the manuscript is accepted for publication.

