## Supplementary material for "Spatial-temporal dynamics of contact among free-ranging domestic dogs *Canis familiaris* in rural Africa"

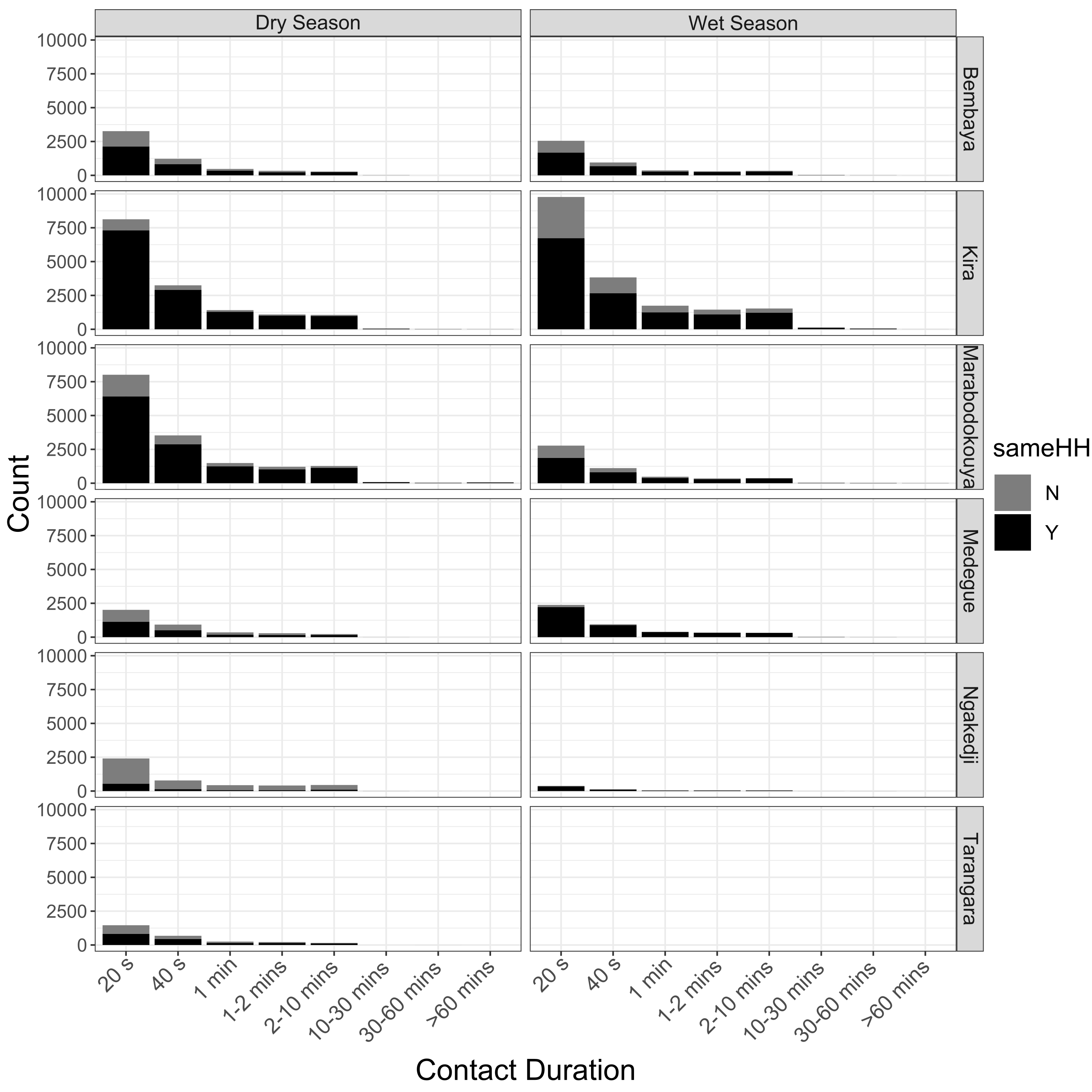

Figure S1: Histograms for the duration of contact events between free-ranging domestic dogs in rural villages in Chad. For each village and season a count of the number of contact events of different durations is plotted for dogs in different households (grey) and in the same household (black). The wet season plot for Tarangara has no bars as no dogs were observed.

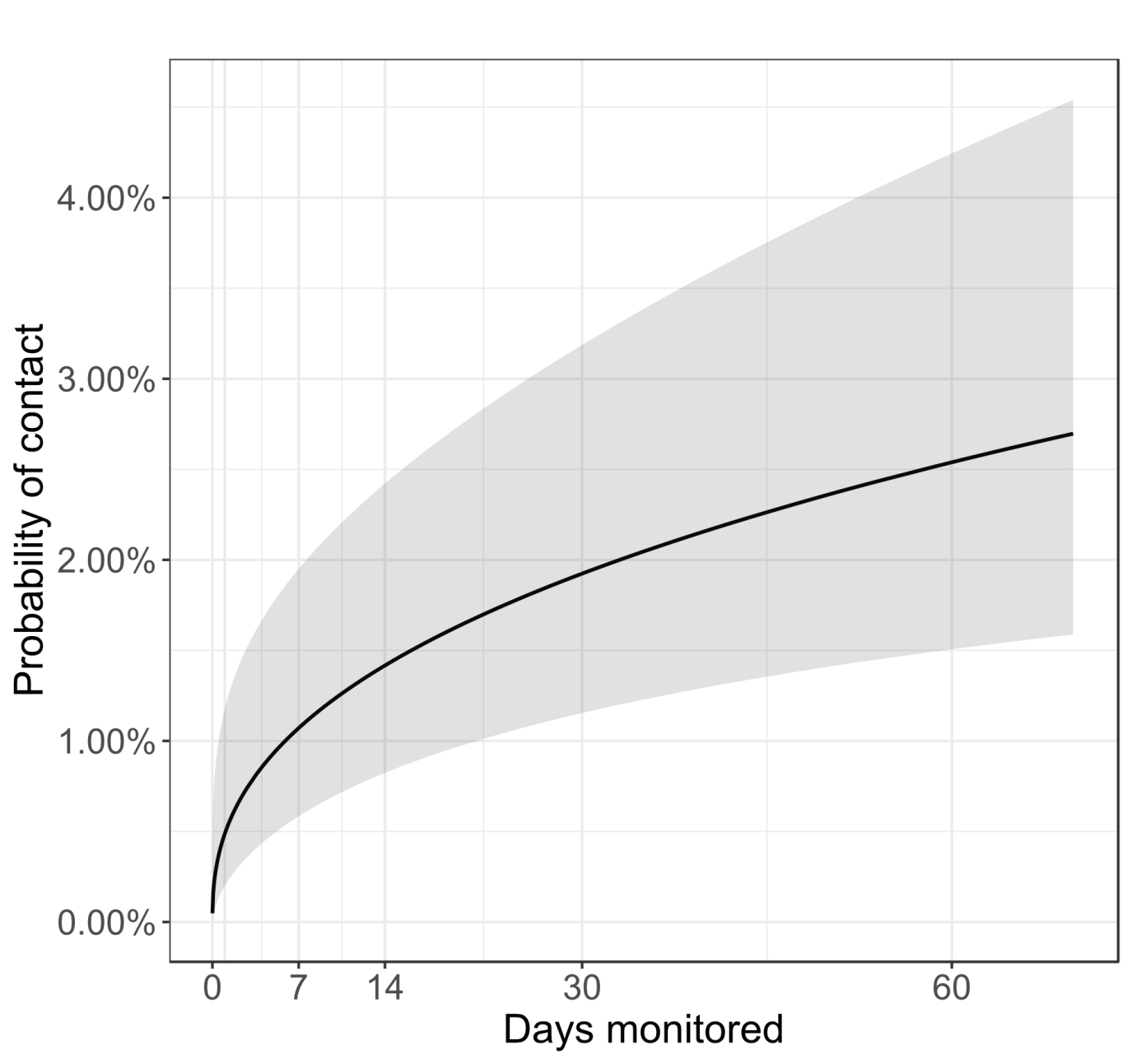

Figure S2: The probability of contact between free-ranging domestic dogs in rural Chad with an increasing number of days over which dyads were observed. The predictions (black line) and confidence interval (shaded area) are plotted from a general linear mixed model.

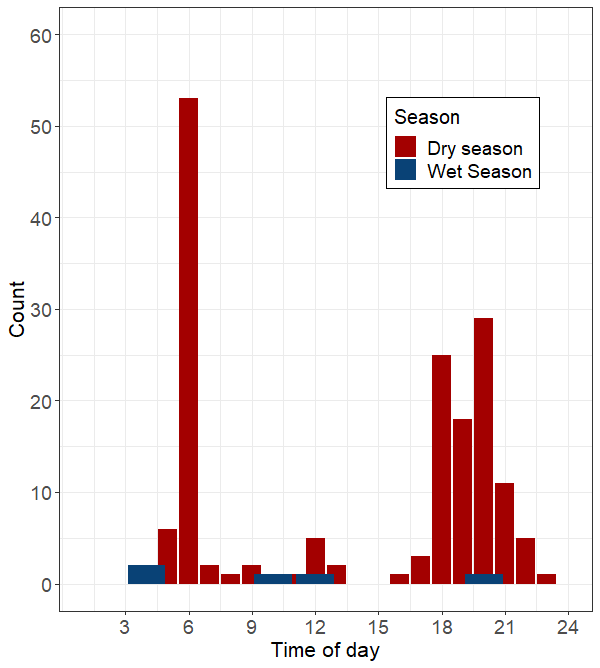

Figure S3: Histogram of contact events for between-village dyads of free-ranging domestic dogs in rural Chad at different times of the day. Counts are presented for contact events in the dry season (red) and wet season (blue).

Table S1. Summary of between-village contact events among free-ranging domestic dogs in rural Chad. For each season and village in the field site ‘Sarh West’, the number of observable dyads is reported with the percentage of the total number of possible dyads in brackets. The number of dyads for which contact was recorded is reported, with the percentage of the number of observable dyads in brackets. The total number of contacts is reported along with the median and inter-quartile range for the duration of contacts. The numbers of contact events in each location category are reported for all spatially referenced contacts.

| Village | Season | No. of observable dyads | No. of dyads in contact | Total no. of contacts | Duration of contacts (s) | Location (around the household : in the village : outside village)* |
| --- | --- | --- | --- | --- | --- | --- |
| Kira | Dry | 1699 (97%) | 14 (1%) | 54 | 20 (20–40) | 1 : 48 : 3 |
|  | Wet | 1250 (61%) | 3 (<1%) | 5 | 20 (20–20) | 0 : 5 : 0 |
| Bembaya | Dry | 838 (97%) | 21 (3%) | 162 | 20 (20–40) | 42 : 98 : 16 |
|  | Wet | 944 (61%) | 3 (<1%) | 5 | 20 (20–20) | 0 : 5 : 0 |
| Ngakedji | Dry | 1489 (97%) | 15 (1%) | 116 | 40 (20–40) | 43 : 56 : 13 |
|  | Wet | 918 (49%) | 0 (0%) | 0 | - | - |
| Overall | Dry | 2013 (97%) | 25 (1%) | 166 | 20 (20–40) | 43 : 101 :16 |
|  | Wet | 1556 (68%) | 3 (<1%) | 5 | 20 (20–20) | 0 : 5 : 0 |
| * Some contacts were not located | | | | | | |

Table S2: Summary of mixed effect models for the probability of free-ranging domestic dogs in rural Chad ever having had contact, their contact probability should they have been in contact and the duration of their contacts. For each model either the odds ratio (OR) or incidence rate ratio (IIR) and confidence intervals (CI) are reported. For the random effects of each model the variance (σ^2^), between-group variance (τ00), group size (Nτ00) and intra-class correlation (ICC) are reported.

|  | **Between Household** | | | | | | | **Within Household** | | | | | |
| --- | --- | --- | --- | --- | --- | --- | --- | --- | --- | --- | --- | --- | --- |
| *Predictors* | **Contact probability** | | **Hourly contact probability** | | | **Contact duration** | | **Contact probability** | | **Hourly contact probability** | | **Contact duration** | |
|  | *OR* | *CI* | *OR* | | *CI* | *IRR* | *CI* | *OR* | *CI* | *OR* | *CI* | *IRR* | *CI* |
| (Intercept) | 6.76 | 0.49 – 92.95 | 0.01 *** | | 0.01 – 0.03 | 99.29 *** | 65.87 – 149.66 | 0.01 | 0.00 – 7.41 | 0.02 *** | 0.01 – 0.03 | 230.93 *** | 190.32 – 280.21 |
| log10(hours monitored) | 2.57 *** | 1.55 – 4.26 | - | | - | - | - | 3.41 | 0.83 – 13.95 | - | - | - | - |
| log2(HH Distance) | 0.36 *** | 0.33 – 0.40 | 0.61 *** | 0.57 – 0.66 | | 0.97 * | 0.93 – 1.00 | - | - | - | - | - | - |
| Sex [female-female] | Reference | | Reference | | | Reference | | Reference | | Reference | | Reference | |
| Sex [male-female] | 1.60 * | 1.10 – 2.34 | 1.17 | 0.92 – 1.49 | | 0.97 | 0.88 – 1.07 | 1.98 | 0.57 – 6.85 | 1.10 | 0.59 – 2.04 | 1.00 | 0.90 – 1.11 |
| Sex [male-male] | 1.26 | 0.70 – 2.30 | 1.22 | 0.85 – 1.74 | | 0.92 | 0.81 – 1.05 | 1.74 | 0.38 – 8.00 | 1.04 | 0.47 – 2.33 | 1.03 | 0.91 – 1.18 |
| Age difference | 1.00 | 0.99 – 1.01 | 1.00 | 0.99 – 1.01 | | 1.01 * | 1.00 – 1.01 | 0.99 | 0.96 – 1.03 | 1.02 * | 1.00 – 1.04 | 1.00 | 0.99 – 1.00 |
| season [dry] | Reference | | Reference | | | Reference | | Reference | | Reference | | Reference | |
| season [wet] | 0.51 | 0.16 – 1.70 | 0.37 ** | 0.20 – 0.72 | | 1.11 | 0.69 – 1.79 | 3.53 | 0.46 – 27.27 | 2.42 *** | 1.91 – 3.07 | 1.08 | 0.85 – 1.37 |
| location [Household] | - | - | Reference | | | Reference | | - | - | Reference | | Reference | |
| location [Out.Village] | - | - | 0.87 | 0.51 – 1.49 | | 1.28 | 0.77 – 2.12 | - | - | 0.10 *** | 0.06 – 0.16 | 1.32 | 0.85 – 2.05 |
| location [Village] | - | - | 1.95 *** | 1.31 – 2.91 | | 1.05 | 0.73 – 1.53 | - | - | 0.91 | 0.74 – 1.13 | 1.08 | 0.87 – 1.34 |
| hour [01] | - | - | Reference | | | Reference | | - | - | Reference | | Reference | |
| hour [02] | - | - | 0.93 | 0.55 – 1.57 | | 0.96 | 0.60 – 1.53 | - | - | 0.77 | 0.60 – 1.00 | 1.01 | 0.79 – 1.29 |
| hour [03] | - | - | 0.75 | 0.43 – 1.29 | | 1.07 | 0.65 – 1.75 | - | - | 0.66 ** | 0.50 – 0.86 | 1.07 | 0.83 – 1.37 |
| hour [04] | - | - | 1.66 * | 1.03 – 2.68 | | 1.34 | 0.88 – 2.04 | - | - | 0.86 | 0.67 – 1.11 | 0.97 | 0.75 – 1.24 |
| hour [05] | - | - | 7.05 *** | 4.74 – 10.50 | | 2.38 *** | 1.68 – 3.38 | - | - | 3.08 *** | 2.50 – 3.79 | 1.79 *** | 1.46 – 2.20 |
| hour [06] | - | - | 6.11 *** | 4.09 – 9.13 | | 1.73 ** | 1.20 – 2.49 | - | - | 2.75 *** | 2.23 – 3.39 | 1.59 *** | 1.30 – 1.94 |
| hour [07] | - | - | 3.57 *** | 2.35 – 5.42 | | 1.33 | 0.91 – 1.93 | - | - | 2.25 *** | 1.82 – 2.79 | 1.30 * | 1.06 – 1.60 |
| hour [08] | - | - | 1.71 * | 1.10 – 2.67 | | 1.20 | 0.82 – 1.78 | - | - | 1.62 *** | 1.30 – 2.03 | 1.21 | 0.98 – 1.50 |
| hour [09] | - | - | 1.10 | 0.69 – 1.77 | | 1.20 | 0.80 – 1.81 | - | - | 1.35 * | 1.07 – 1.70 | 1.24 | 1.00 – 1.55 |
| hour [10] | - | - | 0.73 | 0.44 – 1.19 | | 1.10 | 0.70 – 1.71 | - | - | 1.48 *** | 1.17 – 1.86 | 1.38 ** | 1.11 – 1.71 |
| hour [11] | - | - | 0.70 | 0.42 – 1.15 | | 1.12 | 0.72 – 1.73 | - | - | 1.59 *** | 1.27 – 2.01 | 1.30 * | 1.05 – 1.62 |
| hour [12] | - | - | 1.01 | 0.63 – 1.62 | | 1.44 | 0.96 – 2.16 | - | - | 2.52 *** | 2.03 – 3.13 | 1.50 *** | 1.22 – 1.84 |
| hour [13] | - | - | 1.10 | 0.69 – 1.77 | | 1.20 | 0.80 – 1.81 | - | - | 2.50 *** | 2.01 – 3.10 | 1.62 *** | 1.33 – 1.99 |
| hour [14] | - | - | 1.56 | 0.99 – 2.46 | | 1.30 | 0.87 – 1.95 | - | - | 2.53 *** | 2.04 – 3.14 | 1.54 *** | 1.26 – 1.88 |
| hour [15] | - | - | 2.61 *** | 1.69 – 4.01 | | 1.02 | 0.69 – 1.52 | - | - | 2.59 *** | 2.09 – 3.21 | 1.41 ** | 1.15 – 1.72 |
| hour [16] | - | - | 3.64 *** | 2.40 – 5.52 | | 1.20 | 0.82 – 1.74 | - | - | 2.53 *** | 2.04 – 3.13 | 1.46 *** | 1.19 – 1.79 |
| hour [17] | - | - | 3.81 *** | 2.52 – 5.75 | | 1.29 | 0.89 – 1.87 | - | - | 2.36 *** | 1.91 – 2.92 | 1.21 | 0.99 – 1.49 |
| hour [18] | - | - | 5.54 *** | 3.72 – 8.24 | | 1.15 | 0.80 – 1.65 | - | - | 4.13 *** | 3.37 – 5.05 | 1.59 *** | 1.32 – 1.93 |
| hour [19] | - | - | 6.89 *** | 4.65 – 10.19 | | 1.10 | 0.77 – 1.57 | - | - | 4.30 *** | 3.52 – 5.26 | 1.58 *** | 1.31 – 1.91 |
| hour [20] | - | - | 6.33 *** | 4.27 – 9.39 | | 1.12 | 0.78 – 1.59 | - | - | 2.72 *** | 2.21 – 3.35 | 1.34 ** | 1.09 – 1.63 |
| hour [21] | - | - | 3.28 *** | 2.17 – 4.98 | | 1.35 | 0.93 – 1.95 | - | - | 1.94 *** | 1.56 – 2.41 | 1.21 | 0.99 – 1.50 |
| hour [22] | - | - | 3.48 *** | 2.29 – 5.28 | | 1.24 | 0.85 – 1.80 | - | - | 1.32 * | 1.05 – 1.67 | 1.22 | 0.98 – 1.52 |
| hour [23] | - | - | 2.12 *** | 1.36 – 3.31 | | 1.12 | 0.75 – 1.66 | - | - | 1.09 | 0.86 – 1.39 | 1.03 | 0.81 – 1.29 |
| hour [24] | - | - | 1.40 | 0.87 – 2.26 | | 1.14 | 0.74 – 1.76 | - | - | 0.80 | 0.62 – 1.03 | 1.01 | 0.78 – 1.30 |
| season [wet] * log2(HH Distance) | 1.11 | 0.96 – 1.27 | 1.12 ** | 1.04 – 1.20 | | 1.03 | 0.98 – 1.08 | - | - | - | - | - | - |
| season [wet] * Sex [male-female] | 0.91 | 0.57 – 1.43 | 1.29 * | 1.04 – 1.59 | | 1.04 | 0.90 – 1.20 | 0.46 | 0.05 – 3.82 | 0.53 *** | 0.46 – 0.59 | 1.00 | 0.88 – 1.12 |
| season [wet] * Sex [male-male] | 1.99 * | 1.08 – 3.67 | 1.95 *** | 1.44 – 2.64 | | 1.06 | 0.88 – 1.27 | 0.25 | 0.02 – 3.05 | 1.95 *** | 1.44 – 2.64 | 0.91 | 0.76 – 1.10 |
| season [wet] * Age difference | 0.99 | 0.98 – 1.01 | 1.01 *** | 1.01 – 1.02 | | 1.00 | 1.00 – 1.00 | 1.01 | 0.95 – 1.07 | 0.99 ** | 0.99 – 1.00 | 1.01 ** | 1.00 – 1.01 |
| location [Out.Village] * season [wet] |  |  | 0.38 *** | 0.33 – 0.44 | | 0.63 *** | 0.54 – 0.73 | - | - | 1.21 ** | 1.08 – 1.35 | 0.70 *** | 0.63 – 0.79 |
| location [Village] * season [wet] | - | - | 1.36 *** | 1.22 – 1.51 | | 0.94 | 0.85 – 1.04 | - | - | 2.22 *** | 2.07 – 2.39 | 0.98 | 0.91 – 1.06 |
| hour [02] * season [wet] | - | - | 1.18 | 0.70 – 1.99 | | 0.92 | 0.56 – 1.52 | - | - | 1.06 | 0.79 – 1.42 | 0.98 | 0.73 – 1.31 |
| hour [03] * season [wet] | - | - | 1.07 | 0.64 – 1.78 | | 0.78 | 0.50 – 1.21 | - | - | 1.11 | 0.82 – 1.50 | 0.83 | 0.61 – 1.13 |
| hour [04] * season [wet] | - | - | 0.54 * | 0.33 – 0.90 | | 0.65 | 0.42 – 1.02 | - | - | 0.93 | 0.69 – 1.25 | 0.92 | 0.68 – 1.23 |
| hour [05] * season [wet] | - | - | 0.31 *** | 0.21 – 0.47 | | 0.50 *** | 0.35 – 0.72 | - | - | 0.34 *** | 0.26 – 0.43 | 0.64 *** | 0.50 – 0.82 |
| hour [06] * season [wet] | - | - | 0.30 *** | 0.20 – 0.45 | | 0.77 | 0.54 – 1.11 | - | - | 0.58 *** | 0.45 – 0.73 | 0.76 * | 0.60 – 0.97 |
| hour [07] * season [wet] | - | - | 0.68 | 0.45 – 1.02 | | 0.77 | 0.53 – 1.10 | - | - | 0.73 * | 0.57 – 0.93 | 0.94 | 0.74 – 1.20 |
| hour [08] * season [wet] | - | - | 1.59 * | 1.06 – 2.39 | | 0.67 * | 0.46 – 0.96 | - | - | 0.70 ** | 0.54 – 0.90 | 0.82 | 0.64 – 1.06 |
| hour [09] * season [wet] | - | - | 1.96 ** | 1.28 – 3.00 | | 0.64 * | 0.44 – 0.92 | - | - | 0.61 *** | 0.47 – 0.80 | 0.89 | 0.69 – 1.16 |
| hour [10] * season [wet] | - | - | 3.10 *** | 2.00 – 4.81 | | 0.74 | 0.50 – 1.10 | - | - | 0.48 *** | 0.36 – 0.63 | 0.78 | 0.60 – 1.01 |
| hour [11] * season [wet] | - | - | 2.65 *** | 1.73 – 4.08 | | 0.76 | 0.52 – 1.11 | - | - | 0.35 *** | 0.26 – 0.46 | 0.74 * | 0.57 – 0.96 |
| hour [12] * season [wet] | - | - | 2.70 *** | 1.77 – 4.13 | | 0.47 *** | 0.32 – 0.69 | - | - | 0.28 *** | 0.21 – 0.37 | 0.78 | 0.60 – 1.01 |
| hour [13] * season [wet] | - | - | 2.87 *** | 1.85 – 4.46 | | 0.79 | 0.54 – 1.17 | - | - | 0.28 *** | 0.22 – 0.37 | 0.69 ** | 0.53 – 0.90 |
| hour [14] * season [wet] | - | - | 1.72 * | 1.12 – 2.64 | | 0.68 | 0.46 – 1.00 | - | - | 0.32 *** | 0.25 – 0.43 | 0.71 * | 0.55 – 0.92 |
| hour [15] * season [wet] | - | - | 0.90 | 0.59 – 1.36 | | 0.82 | 0.56 – 1.20 | - | - | 0.40 *** | 0.30 – 0.52 | 0.75 * | 0.58 – 0.96 |
| hour [16] * season [wet] | - | - | 0.74 | 0.49 – 1.11 | | 0.74 | 0.51 – 1.06 | - | - | 0.45 *** | 0.35 – 0.58 | 0.71 ** | 0.56 – 0.91 |
| hour [17] * season [wet] | - | - | 0.85 | 0.57 – 1.26 | | 0.65 * | 0.45 – 0.93 | - | - | 0.71 ** | 0.55 – 0.91 | 0.90 | 0.71 – 1.15 |
| hour [18] * season [wet] | - | - | 1.04 | 0.71 – 1.53 | | 0.87 | 0.61 – 1.23 | - | - | 0.92 | 0.72 – 1.17 | 0.85 | 0.67 – 1.07 |
| hour [19] * season [wet] | - | - | 0.89 | 0.61 – 1.29 | | 0.82 | 0.58 – 1.15 | - | - | 0.91 | 0.72 – 1.16 | 0.83 | 0.66 – 1.05 |
| hour [20] * season [wet] | - | - | 0.86 | 0.59 – 1.25 | | 0.78 | 0.56 – 1.10 | - | - | 1.05 | 0.82 – 1.35 | 0.94 | 0.74 – 1.19 |
| hour [21] * season [wet] | - | - | 0.90 | 0.61 – 1.33 | | 0.81 | 0.56 – 1.15 | - | - | 1.00 | 0.78 – 1.30 | 1.00 | 0.78 – 1.28 |
| hour [22] * season [wet] | - | - | 0.75 | 0.50 – 1.12 | | 0.88 | 0.61 – 1.26 | - | - | 1.00 | 0.76 – 1.31 | 0.90 | 0.69 – 1.17 |
| hour [23] * season [wet] | - | - | 0.97 | 0.64 – 1.49 | | 0.83 | 0.56 – 1.22 | - | - | 1.05 | 0.79 – 1.38 | 0.96 | 0.73 – 1.27 |
| hour [24] * season [wet] | - | - | 1.05 | 0.67 – 1.65 | | 0.91 | 0.61 – 1.38 | - | - | 1.13 | 0.85 – 1.51 | 0.99 | 0.75 – 1.32 |
| hour [02] * location [Out.Village] | - | - | 0.39 * | 0.16 – 0.97 | | 1.00 | 0.43 – 2.29 | - | - | 0.73 | 0.34 – 1.57 | 1.87 | 0.92 – 3.79 |
| hour[02] * location [Village] | - | - | 0.65 | 0.36 – 1.15 | | 1.09 | 0.62 – 1.89 | - | - | 1.24 | 0.91 – 1.68 | 0.97 | 0.72 – 1.33 |
| hour[03] * location [Out.Village] | - | - | 0.61 | 0.25 – 1.49 | | 1.72 | 0.81 – 3.68 | - | - | 1.32 | 0.66 – 2.61 | 2.91 *** | 1.59 – 5.34 |
| hour[03] * location [Village] | - | - | 1.16 | 0.64 – 2.09 | | 0.95 | 0.56 – 1.63 | - | - | 1.19 | 0.86 – 1.63 | 1.07 | 0.77 – 1.47 |
| hour[4] * location [Out.Village] | - | - | 0.32 ** | 0.14 – 0.75 | | 1.49 | 0.74 – 2.98 | - | - | 0.59 | 0.26 – 1.31 | 1.67 | 0.79 – 3.53 |
| hour[4] * location [Village] | - | - | 0.66 | 0.38 – 1.14 | | 1.15 | 0.70 – 1.89 | - | - | 1.10 | 0.81 – 1.49 | 1.13 | 0.83 – 1.55 |
| hour[5] * location [Out.Village] | - | - | 0.41 ** | 0.22 – 0.76 | | 0.76 | 0.43 – 1.34 | - | - | 3.91 *** | 2.35 – 6.50 | 1.12 | 0.70 – 1.81 |
| hour[5] * location [Village] | - | - | 0.73 | 0.47 – 1.13 | | 0.85 | 0.57 – 1.27 | - | - | 1.47 ** | 1.14 – 1.89 | 0.91 | 0.70 – 1.18 |
| hour[6] * location [Out.Village] | - | - | 0.90 | 0.50 – 1.62 | | 1.15 | 0.66 – 2.00 | - | - | 4.40 *** | 2.66 – 7.29 | 0.99 | 0.62 – 1.60 |
| lhour[6] * ocation [Village] | - | - | 0.74 | 0.47 – 1.16 | | 0.95 | 0.63 – 1.44 | - | - | 1.21 | 0.93 – 1.56 | 0.96 | 0.74 – 1.24 |
| hour[7] * location [Out.Village] | - | - | 1.18 | 0.64 – 2.15 | | 1.20 | 0.69 – 2.10 | - | - | 6.11 *** | 3.70 – 10.09 | 1.07 | 0.67 – 1.72 |
| hour[7] * location [Village] | - | - | 0.75 | 0.47 – 1.19 | | 0.96 | 0.63 – 1.47 | - | - | 1.25 | 0.96 – 1.62 | 0.92 | 0.71 – 1.19 |
| hour[8] * location [Out.Village] | - | - | 2.36 ** | 1.28 – 4.34 | | 2.05 * | 1.17 – 3.61 | - | - | 9.31 *** | 5.62 – 15.43 | 1.17 | 0.73 – 1.88 |
| hour[8] * location [Village] | - | - | 0.88 | 0.55 – 1.43 | | 1.04 | 0.67 – 1.62 | - | - | 1.07 | 0.82 – 1.41 | 0.99 | 0.76 – 1.31 |
| hour[9] * location [Out.Village] | - | - | 2.75 ** | 1.46 – 5.17 | | 1.66 | 0.94 – 2.94 | - | - | 9.43 *** | 5.65 – 15.74 | 1.07 | 0.66 – 1.73 |
| hour[9] * location [Village] | - | - | 0.91 | 0.55 – 1.51 | | 1.02 | 0.64 – 1.61 | - | - | 1.12 | 0.84 – 1.49 | 0.91 | 0.68 – 1.21 |
| hour[10] * location [Out.Village] | - | - | 3.61 *** | 1.91 – 6.83 | | 1.30 | 0.72 – 2.34 | - | - | 7.79 *** | 4.65 – 13.05 | 0.89 | 0.55 – 1.43 |
| hour[10] * location [Village] | - | - | 0.90 | 0.54 – 1.51 | | 1.04 | 0.65 – 1.68 | - | - | 1.03 | 0.77 – 1.37 | 0.90 | 0.68 – 1.19 |
| hour[11] * location [Out.Village] | - | - | 3.96 *** | 2.07 – 7.55 | | 1.23 | 0.68 – 2.24 | - | - | 5.59 *** | 3.31 – 9.44 | 1.04 | 0.64 – 1.70 |
| hour[11] * location [Village] | - | - | 1.35 | 0.80 – 2.27 | | 1.05 | 0.65 – 1.69 | - | - | 1.31 | 0.98 – 1.73 | 0.98 | 0.74 – 1.29 |
| hour[12] * location [Out.Village] | - | - | 2.62 ** | 1.40 – 4.89 | | 1.73 | 0.97 – 3.09 | - | - | 2.72 *** | 1.60 – 4.62 | 1.05 | 0.64 – 1.71 |
| hour[12] * location [Village] | - | - | 0.98 | 0.60 – 1.60 | | 1.16 | 0.73 – 1.83 | - | - | 0.98 | 0.74 – 1.28 | 0.88 | 0.67 – 1.15 |
| hour[13] * location [Out.Village] | - | - | 1.28 | 0.67 – 2.46 | | 1.25 | 0.69 – 2.25 | - | - | 2.21 ** | 1.29 – 3.79 | 0.89 | 0.54 – 1.46 |
| hour[13] * location [Village] | - | - | 0.74 | 0.45 – 1.20 | | 0.83 | 0.53 – 1.28 | - | - | 1.03 | 0.79 – 1.35 | 0.89 | 0.69 – 1.17 |
| hour[14] * location [Out.Village] | - | - | 0.92 | 0.47 – 1.78 | | 1.32 | 0.74 – 2.38 | - | - | 1.43 | 0.82 – 2.51 | 1.71 * | 1.03 – 2.84 |
| hour[14] * location [Village] | - | - | 0.82 | 0.50 – 1.33 | | 0.89 | 0.57 – 1.38 | - | - | 0.90 | 0.69 – 1.18 | 0.94 | 0.72 – 1.23 |
| hour[15] * location [Out.Village] | - | - | 0.92 | 0.49 – 1.74 | | 0.83 | 0.46 – 1.51 | - | - | 1.46 | 0.84 – 2.54 | 0.99 | 0.59 – 1.67 |
| hour[15] * location [Village] | - | - | 0.77 | 0.48 – 1.23 | | 1.02 | 0.66 – 1.58 | - | - | 0.96 | 0.74 – 1.25 | 0.98 | 0.75 – 1.27 |
| hour[16] * location [Out.Village] | - | - | 1.03 | 0.56 – 1.89 | | 1.15 | 0.66 – 2.03 | - | - | 2.54 *** | 1.50 – 4.30 | 0.84 | 0.51 – 1.38 |
| hour[16] * location [Village] | - | - | 0.73 | 0.46 – 1.16 | | 0.94 | 0.62 – 1.44 | - | - | 1.06 | 0.81 – 1.37 | 0.90 | 0.69 – 1.17 |
| hour[17] * location [Out.Village] | - | - | 0.73 | 0.39 – 1.34 | | 1.18 | 0.67 – 2.07 | - | - | 2.13 ** | 1.26 – 3.61 | 1.15 | 0.70 – 1.88 |
| hour[17] * location [Village] | - | - | 0.79 | 0.50 – 1.25 | | 0.93 | 0.61 – 1.41 | - | - | 1.00 | 0.77 – 1.30 | 1.02 | 0.79 – 1.32 |
| hour[18] * location [Out.Village] | - | - | 0.42 ** | 0.23 – 0.77 | | 0.65 | 0.36 – 1.18 | - | - | 0.88 | 0.52 – 1.50 | 1.12 | 0.68 – 1.85 |
| hour[18] * location [Village] | - | - | 0.78 | 0.51 – 1.21 | | 0.91 | 0.61 – 1.36 | - | - | 0.77 * | 0.60 – 0.98 | 0.85 | 0.67 – 1.08 |
| hour[19] * location [Out.Village] | - | - | 0.38 ** | 0.21 – 0.69 | | 0.72 | 0.41 – 1.28 | - | - | 0.46 ** | 0.26 – 0.81 | 0.87 | 0.50 – 1.51 |
| hour[19] * location [Village] | - | - | 0.97 | 0.63 – 1.48 | | 1.03 | 0.69 – 1.53 | - | - | 0.75 * | 0.59 – 0.96 | 0.90 | 0.71 – 1.15 |
| hour[20] * location [Out.Village] | - | - | 0.52 * | 0.29 – 0.94 | | 0.80 | 0.46 – 1.40 | - | - | 0.56 | 0.31 – 1.01 | 0.85 | 0.47 – 1.51 |
| hour[20] * location [Village] | - | - | 0.92 | 0.60 – 1.42 | | 0.99 | 0.67 – 1.48 | - | - | 0.81 | 0.63 – 1.04 | 0.87 | 0.68 – 1.12 |
| hour[21] * location [Out.Village] | - | - | 1.03 | 0.56 – 1.89 | | 0.54 * | 0.30 – 0.96 | - | - | 0.84 | 0.47 – 1.52 | 0.82 | 0.47 – 1.41 |
| hour[21] * location [Village] | - | - | 1.23 | 0.78 – 1.93 | | 0.80 | 0.53 – 1.22 | - | - | 0.97 | 0.75 – 1.27 | 0.90 | 0.70 – 1.17 |
| hour[22] * location [Out.Village] | - | - | 0.71 | 0.38 – 1.32 | | 0.77 | 0.43 – 1.38 | - | - | 1.31 | 0.73 – 2.37 | 0.77 | 0.44 – 1.34 |
| hour[22] * location [Village] | - | - | 0.89 | 0.57 – 1.42 | | 0.88 | 0.58 – 1.34 | - | - | 0.97 | 0.73 – 1.28 | 0.91 | 0.69 – 1.20 |
| hour[23] * location [Out.Village] | - | - | 0.80 | 0.41 – 1.55 | | 1.01 | 0.55 – 1.85 | - | - | 1.19 | 0.64 – 2.23 | 0.95 | 0.53 – 1.71 |
| hour[23] * location [Village] | - | - | 0.84 | 0.52 – 1.36 | | 0.98 | 0.63 – 1.54 | - | - | 1.02 | 0.76 – 1.36 | 1.01 | 0.76 – 1.35 |
| hour[24] * location [Out.Village] | - | - | 0.77 | 0.38 – 1.59 | | 0.87 | 0.45 – 1.67 | - | - | 1.21 | 0.62 – 2.33 | 1.15 | 0.63 – 2.10 |
| hour[24] * location [Village] | - | - | 0.93 | 0.56 – 1.56 | | 0.89 | 0.55 – 1.45 | - | - | 1.27 | 0.94 – 1.71 | 1.01 | 0.75 – 1.36 |
|  | **Random Effects** | | | | | | | | | | | | |
| σ^2^ | 3.29 | | 3.29 | | | 0.79 | | 3.29 | | 3.29 | | 1.76 | |
| τ_00_ [ID1] | 1.16 | | 0.18 | | | < 0.01 | | < 0.01 | | 0.48 | | 0.01 | |
| τ00 [ID2] | 0.87 | | 0.10 | | | < 0.01 | | < 0.01 | | 0.36 | | < 0.01 | |
| τ00 [Household ID1] | 0.96 | | 0.47 | | | 0.02 | | 0.37 | | 0.3 | | < 0.01 | |
| τ00 [Household ID2] | 0.27 | | 0.15 | | | 0.05 | | 0.02 | | 0.18 | | < 0.01 | |
| τ00 [Dyad ID] | - | | 0.82 | | | 0.01 | | - | | 0.61 | | < 0.01 | |
| ICC | 0.5 | | 0.34 | | |  | | 0.5 | | 0.37 | | 0.02 | |
| Nτ00 [ID1] | 251 | | 182 | | | 186 | | 89 | | 71 | | 90 | |
| Nτ00 [ID2] | 257 | | 191 | | | 182 | | 92 | | 74 | | 90 | |
| Nτ00 [Household ID1] | 156 | | 125 | | | 128 | | 63 | | 50 | | 48 | |
| Nτ00 [Household ID2] | 160 | | 131 | | | 126 | | 63 | | 50 | | 48 | |
| Nτ00 [Dyad ID] | - | | 953 | | | 956 | | - | | 105 | | 104 | |
| Observations | 7022 | | 85164 | | | 6985 | | 168 | | 9492 | | 13609 | |
| Marginal R^2^ / Conditional R^2^ | 0.424 / 0.711 | | 0.201 / 0.476 | | | 0.074 / 0.152 | | 0.069 / 0.109 | | 0.198 / 0.494 | | 0.017 / 0.031 | |
| * p<0.05   ** p<0.01   *** p<0.001 | | | | | | | | | | | | | |

| *Hour* | **Hour: hour[i] - hour[i+3]** | | **Hour*season:  hour[Dry] - hour[Wet]** | | **Hour*location:  hour[in village] - hour [outside]** | |
| --- | --- | --- | --- | --- | --- | --- |
|  | *Odds Ratios* | *95% CI* | *Odds Ratios* | *95% CI* | *Odds Ratios* | *95% CI* |
| 01:00 | 1.37 | 0.76 - 2.45 | 0.77 | 0.53 - 1.10 | 4.23*** | 2.39 - 7.49 |
| 02:00 | 0.24*** | 0.14 - 0.42 | 0.65* | 0.43 - 0.99 | 7.01*** | 2.94 - 16.74 |
| 03:00 | 0.24*** | 0.14 - 0.39 | 0.71 | 0.48 - 1.07 | 8.04*** | 3.67 - 17.59 |
| 04:00 | 0.26*** | 0.16 - 0.43 | 1.41 | 0.95 - 2.10 | 8.66*** | 3.98 - 18.83 |
| 05:00 | 0.95 | 0.69 - 1.31 | 2.46*** | 1.90 - 3.18 | 7.43*** | 5.16 - 10.71 |
| 06:00 | 1.39* | 1.00 - 1.93 | 2.54*** | 1.96 - 3.31 | 3.48*** | 2.58 - 4.69 |
| 07:00 | 1.49** | 1.07 - 2.08 | 1.12 | 0.87 - 1.45 | 2.70*** | 1.96 - 3.73 |
| 08:00 | 1.39* | 1.00 - 1.93 | 0.48*** | 0.37 - 0.62 | 1.59** | 1.16 - 2.16 |
| 09:00 | 0.92 | 0.66 - 1.29 | 0.39*** | 0.29 - 0.52 | 1.40 | 0.99 - 1.98 |
| 10:00 | 1.03 | 0.71 - 1.49 | 0.25*** | 0.18 - 0.33 | 1.06 | 0.75 - 1.50 |
| 11:00 | 1.07 | 0.74 - 1.54 | 0.29*** | 0.22 - 0.39 | 1.44* | 1.03 - 2.02 |
| 12:00 | 1.03 | 0.74 - 1.44 | 0.28*** | 0.21 - 0.37 | 1.59** | 1.14 - 2.21 |
| 13:00 | 0.64*** | 0.46 - 0.90 | 0.27*** | 0.20 - 0.36 | 2.43*** | 1.61 - 3.67 |
| 14:00 | 0.64*** | 0.46 - 0.90 | 0.44*** | 0.33 - 0.59 | 3.77*** | 2.43 - 5.84 |
| 15:00 | 0.57*** | 0.41 - 0.77 | 0.85 | 0.65 - 1.12 | 3.51*** | 2.38 - 5.17 |
| 16:00 | 0.62*** | 0.46 - 0.82 | 1.03 | 0.80 - 1.33 | 3.02*** | 2.17 - 4.20 |
| 17:00 | 0.63*** | 0.48 - 0.84 | 0.90 | 0.71 - 1.15 | 4.62*** | 3.24 - 6.59 |
| 18:00 | 1.15 | 0.87 - 1.53 | 0.73* | 0.59 - 0.91 | 7.97*** | 5.58 - 11.38 |
| 19:00 | 1.80*** | 1.33 - 2.43 | 0.86 | 0.70 - 1.05 | 10.82*** | 7.71 - 15.18 |
| 20:00 | 2.51*** | 1.82 - 3.48 | 0.89 | 0.72 - 1.09 | 7.53*** | 5.51 - 10.29 |
| 21:00 | 2.62*** | 1.79 - 3.83 | 0.85 | 0.67 - 1.07 | 5.05*** | 3.67 - 6.93 |
| 22:00 | 2.58*** | 1.69 - 3.95 | 1.02 | 0.79 - 1.32 | 5.37*** | 3.68 - 7.83 |
| 23:00 | 2.87*** | 1.65 - 5.01 | 0.79 | 0.59 - 1.05 | 4.44*** | 2.85 - 6.92 |
| 00:00 | 1.87*** | 1.06 - 3.29 | 0.73 | 0.53 - 1.01 | 5.10*** | 2.98 - 8.72 |
| * p<0.05   ** p<0.01   *** p<0.001 | | | | | | |

Table S3: Contrasts for model predictions of the hourly probability that free-ranging domestic dogs in rural Chad were in contact. For each hour of the day the odds ratio and 95% confidence interval are provided for the contrast between the probability of contact at the focal hour and to that three hours ahead, between the focal hour in the wet and dry season, and between contacts at the focal hour around the village (within 100 m of a household with tracked dogs) and outside of the village
